# Gene loss and acquisition in lineages of bacteria evolving in a human host environment

**DOI:** 10.1101/2020.02.03.931667

**Authors:** Migle Gabrielaite, Helle K. Johansen, Søren Molin, Finn C. Nielsen, Rasmus L. Marvig

**Author notes:** To whom correspondence should be addressed; Migle Gabrielaite, Rasmus L. Marvig.

## Abstract

While genome analyses have documented that there are differences in the gene repertoire between evolutionary distant lineages of the same bacterial species, less is known about micro-evolutionary dynamics of gene loss and acquisition within lineages of bacteria as they evolve over the timescale of years. This knowledge is valuable to understand both the basic mutational steps that on long timescales lead to evolutionary distant bacterial lineages, and the evolution of the individual lineages themselves. In the case that lineages evolve in a human host environment, gene loss and acquisition may furthermore have implication for disease.

We analyzed the genomes of 45 *Pseudomonas aeruginosa* lineages evolving in the lungs of cystic fibrosis patients to identify genes that are lost or acquired during the first years of infection in each of the different lineages. On average, the lineage genome content changed with 88 genes (range 0–473). Genes were more often lost than acquired, and prophage genes were more variable than bacterial genes. We identified genes that were lost or acquired independently across different clonal lineages, i.e. convergent molecular evolution. Convergent evolution suggests that there is a selection for loss and acquisition of certain genes in the host environment. We find that a significant proportion of such genes are associated with virulence; a trait previously shown to be important for adaptation. Furthermore, we also compared the genomes across lineages to show that within-lineage variable genes more often belonged to genomic content not shared across all lineages. Finally, we used 4,760 genes shared by 446 *P. aeruginosa* genomes to develop a stable and discriminatory typing scheme for *P. aeruginosa* clone types (Pactyper, https://github.com/MigleSur/Pactyper). In sum, our analysis adds to the knowledge on the pace and drivers of gene loss and acquisition in bacteria evolving over multiple years in a human host environment and provides a basis to further understand how gene loss and acquisition plays a role in lineage differentiation and host adaptation.

**Data Summary:** *P. aeruginosa* genome sequencing data has been made publicly available by Marvig *et al.* (2015) and is deposited in Sequence Read Archive (SRA) under accession ERP004853.

## Introduction

Gene acquisition and gene loss are prominent in bacterial evolution and are also important for adaptation. [1, 2] In contrast to point mutations, small insertions and deletions (microindels), inversions, and translocations that gradually alter existing genomic content, the acquisition or loss of entire genes rapidly confers large changes to the genomic content, and gene acquisition and loss alters phenotypes such as virulence, antibiotic resistance, and metabolic capability. [3, 4] Thus, genome-wide analysis of the gene presence or absence is necessary to better understand the bacterial evolution and adaptation.

While genome comparison of evolutionary distant lineages of the same bacterial species gives insight of gene flux over macro-evolutionary scales, there is much less knowledge of the pace and mechanisms by which genes are lost and acquired at micro-evolutionary scales, i.e. from studies of evolution of individual bacterial lineages. Additionally, we only have a limited understanding of how lineage gene loss and acquisition is driven by selective versus genetic drift. [1, 2]

Evolutionary studies on individual bacterial lineages are dependent on the ability to obtain multiple samples of the same lineage which can be difficult in natural, *in vivo*, environments that constantly change, so studies are more easily performed *in vitro*. However, *P. aeruginosa* infections in cystic fibrosis (CF) patients represent an infectious disease scenario in which the genomic evolution of individual bacterial lineages can be followed over years; thus, gives an opportunity to research bacterial evolution and adaptation *in vivo* in the human host. [5, 6] Already there is large knowledge on the role of point mutations and microindels in evolution and adaptation of *P. aeruginosa* in CF patients, whereas gene loss and acquisition has been less investigated. A better understanding about the genetic changes responsible for *P. aeruginosa* pathogenicity in CF patients is crucial to improve the CF treatment strategies. [7, 8, 9]

To better understand the role of gene loss and acquisition for within-host evolution and adaptation, we used genomic data from 474 longitudinally collected isolates of *P. aeruginosa* from children and young CF patients to investigate gene loss and acquisition in clonal lineages of *P. aeruginosa* as they evolve from initial invasion of cystic fibrosis airways and onwards as they adapt to the human host. In total 34 patients and 45 different clonal lineages were analyzed, and we aimed to identify gene loss or acquisition events in each of the different lineages, and to identify patterns of gene loss and acquisition across lineages to better understand the genetic basis of bacterial adaptation in the human host.

## Materials and methods

### Bacterial isolates, determination of clone types, and lineage definition

The study used genomic data from a previously reported collection of 474 clinical isolates of *P. aeruginosa* that were sampled from 34 patients with CF attending the Copenhagen Cystic Fibrosis Center at the University Hospital, Rigshospitalet, Denmark [10]. Genomes were sequenced as follows: genomic DNA was prepared from *P. aeruginosa* isolates on a QIAcube system using a DNeasy Blood and Tissue kit (Qiagen) and sequenced on an Illumina HiSeq 2000 platform, generating 100-bp paired-end reads and using a multiplexed protocol to obtain an average of 7,139,922 reads (range of 3,111,062–13,085,190) for each of the genomic libraries.

The clone type of each of the isolates was previously reported by Marvig *et al.,* (2015) that determined the clone types by an *ad hoc* analysis [10], and we also confirmed the clone types by Pactyper (https://github.com/MigleSur/Pactyper) which is a tool developed as part of this study for stable and discriminatory clone typing of bacterial genomes. Pactyper was run with default settings which defined isolates to be of different clone types if they are different by more than 5,000 SNPs in a core genome of 4,760 genes (i.e. all genes shared by 446 of 474 genomes that were successfully *de novo* assembled (see below)). Isolates of the same clone type and from the same patient were defined as being part of the same lineage.

### Bacterial genome assembly

Sequence reads from each isolate were error corrected and *de novo* assembled by SPAdes version 3.10.1 [11] using k-mer sizes from 21 to 127. Various sizes of k-mers are used for different fragments, thus non-uniform coverage problem is overcome. Assembled contigs were joined to scaffolds as per SPAdes default parameters. *De novo* assemblies of sequence reads from 28 of the 474 isolates (6%) were unsuccessful (more than 500 scaffolds in the final assembly), and the isolates were excluded from the analysis.

### Genome annotation and identification of gene loss and acquisition within lineage pan-genomes

*De novo* assembled genomes were annotated using Prokka version 1.11 [14] with the settings of a minimum contig length of 200 nucleotides and using custom annotation database for *P. aeruginosa* species based on PAO1 (RefSeq assembly accession: GCF_000006765.1) and UCBPP-PA14 (RefSeq assembly accession: GCF_000014625.1) reference genomes. GenAPI was run with default settings to determine lineage pan-genomes, and the presence and absence of genes in individual genomes. [15] Note, GenAPI default settings include that genes shorter than 150 nucleotides were excluded from the analysis.

### Aggregated pan-genome and visualization

An aggregated pan-genome was determined by clustering all genes in the lineage pan-genomes with CD-HIT-EST version 4.6.1 software [18] and the requirement for alignments to cover at least 80% of the query gene length and to have minimum of 90% identity of the alignment. These thresholds are equivalent to default thresholds of GenAPI which was used to determine lineage pan-genomes. Every gene in the aggregated pan-genome was then aligned back to the individual lineage pan-genomes to determine if the gene is (1) non-present in the lineage pan-genome, (2) present and variable within the lineage or (3) present and non-variable within the lineage. A heatmap for the aggregated pan-genome was made with all 45 lineages using R version 3.3.3 [16] and pheatmap library version 1.0.8 [17].

### Identification of most variable genes

For the gene to be considered highly variable, it had to be lost or acquired (variable) in at least 4 lineages. All genes which were identified as highly variable were manually inspected as follows to confirm the results: Read sequences from isolates of interest were aligned to the reference PAO1 (RefSeq assembly accession: GCF_000006765.1) genome using bowtie2 version 2.3.2 [12] with the default parameters for paired-end sequencing. The sequence alignments at genomic positions of interest were visualized with IGV version 2.4.9 [13] and then manually assessed. Non-*Pseudomonas* origin genes were manually inspected by evaluating their alignments to the pan-genome genes (from GenAPI analysis).

In total, two genes (PA1352 and PA3457) were concluded to be falsely called as variable because the alignments to the reference PAO1 genome did not support the prediction of the genes being absent. These false calls were in genome regions which are complex and difficult to *de novo* assemble, i.e. calls of gene presence or absence showed to vary with the success of the assembly of the specific genome region rather than the result of genuine gene presence or absence.

### Reanalysis of *P. aeruginosa* genomes previously analyzed in other *P. aeruginosa* study

Analysis of the dataset from Freschi *et al.* (2019) study [19] included 1,139 out of 1,311 genomes as 172 genomes were not publicly available on the day of access (2019-04-02). All available samples were analyzed with SaturnV (https://github.com/ejfresch/saturnV) using the default settings and the lazy option, and with GenAPI (https://github.com/MigleSur/GenAPI) [15] using default settings.

### Resistance, virulence, pathoadaptive and prophage origin gene identification

Resistance, plasmid and virulence genes were identified by comparing the aggregated pan-genome of 45 lineages with the corresponding databases by using ABRicate version 0.8. [20] The gene was considered present from the corresponding database if its identity was at least 98% and the alignment made up minimum 25% of the gene length.

PlasmidFinder [21] database (263 sequences; retrieved: 2018-03-21) was used for plasmid gene, VFDB (2,597 genes; retrieved: 2018-03-21) [22]—for virulence gene and Resfinder (2,280 genes; retrieved: 2018-03-21) [23]—for resistance gene identification, ACLAME database (54,945 genes; 2018-06-07) [24]—for prophage origin sequence identification. For pathoadaptive gene identification, 52 pathoadaptive gene list from Marvig *et al.* (2015) [10] was compared to the aggregated pan-genome of the 45 lineages.

Fisher’s exact test was performed to identify whether the number of genes from the corresponding database is significantly different between within-host variable and non-variable genes.

### Genomic island identification

Genomic islands were predicted using IslandViewer 4 [25] webserver with PAO1 (RefSeq assembly accession: GCF_000006765.1) as reference genome. Genomic island prediction was performed on the annotated scaffold sequences. IslandViewer 4 integrated tools—IslandPath-DIMOB and SIGI-HMM— were used for prediction of genomic islands.

### Pairwise gene and SNP distance estimation between *P. aeruginosa* isolates

Gene distance between genomes was defined as the number of genes not present in one of the genomes as determined by GenAPI (i.e.genes present in one genome and absent in the other, and *vice versa*). Pairwise SNP distance was determined using PacTyper (https://github.com/MigleSur/Pactyper) that uses sequence reads to call and compare SNPs across the core genome. Default thresholds of Pactyper requires that sequence reads cover at least 80% of the core genome with not less than 10-fold coverage to ensure exclusion of genomes with poor sequencing coverage. The core genome was defined in this study with GenAPI analysis by including all genes shared by the 446 successfully sequenced *P. aeruginosa* genomes. The core genome contained 4,760 genes (4,705,617 nucleotides).

### Phylogenetic tree generation

A SNP-based phylogenetic tree was generated with RAxML version 8.2.11 [26] (with the GTRCAT settings for nucleotide sequence analysis and 12345 as a random number seed) by alignments of the previously defined core genome, and PAO1 (RefSeq assembly accession: GCF_000006765.1) and UCBPP-PA14 (RefSeq assembly accession: GCF_000014625.1) were included as reference genomes. A gene presence-absence based phylogenetic tree was generated with RAxML version 8.2.11 [26] (with the BINCAT settings for nucleotide sequence analysis and 12345 as a random number seed) by using gene presence-absence information from GenAPI analysis, and PAO1 (RefSeq assembly accession: GCF_000006765.1) and UCBPP-PA14 (RefSeq assembly accession: GCF_000014625.1) were included as reference genomes. Microreact webservice was used to visualize the phylogenetic trees. [27]

## Results

### *De novo* genome assembly and gene annotation

We have previously generated short read sequence data for the genomes of 474 isolates of *P. aeruginosa* sampled from the airways of 34 young CF patients to follow the genomic evolution of bacterial lineages within the host airways over the initial 0–9 years of infection. [10] While the previous analysis aligned sequence reads to a *P. aeruginosa* reference genome to identify single nucleotide polymorphisms (SNPs) and small insertions and deletions (indels), we here use the same sequence reads, but for *de novo* assembly of genomes, to identify genes that are either lost or acquired during the course of infection.

We were successful to *de novo* assemble the genomes of 446 isolates into 500 scaffolds or less (median 172 scaffolds). Sizes of the assembled genomes ranged over 6,032,338–7,593,423 nucleotides (nt), and they contained 5,462–7,111 genes. The 446 assembled genomes represented 51 clone types as per defined by Marvig *et al.* (2015) [10]. We grouped the isolates into 45 lineages, i.e. isolates of the same clone type and from the same patient were grouped together, to allow identification of within-host accumulated gene differences (Figure 1). In total, the 45 lineages encompassed 423 isolates distributed on 34 patients and 34 clone types. The remainder 23 isolates with successful genome assembly were excluded from the analysis as there were no other clonal genomes available for the respective patients, i.e. at least two genomes were required for intra-lineage genome comparison.

**Figure 1.**
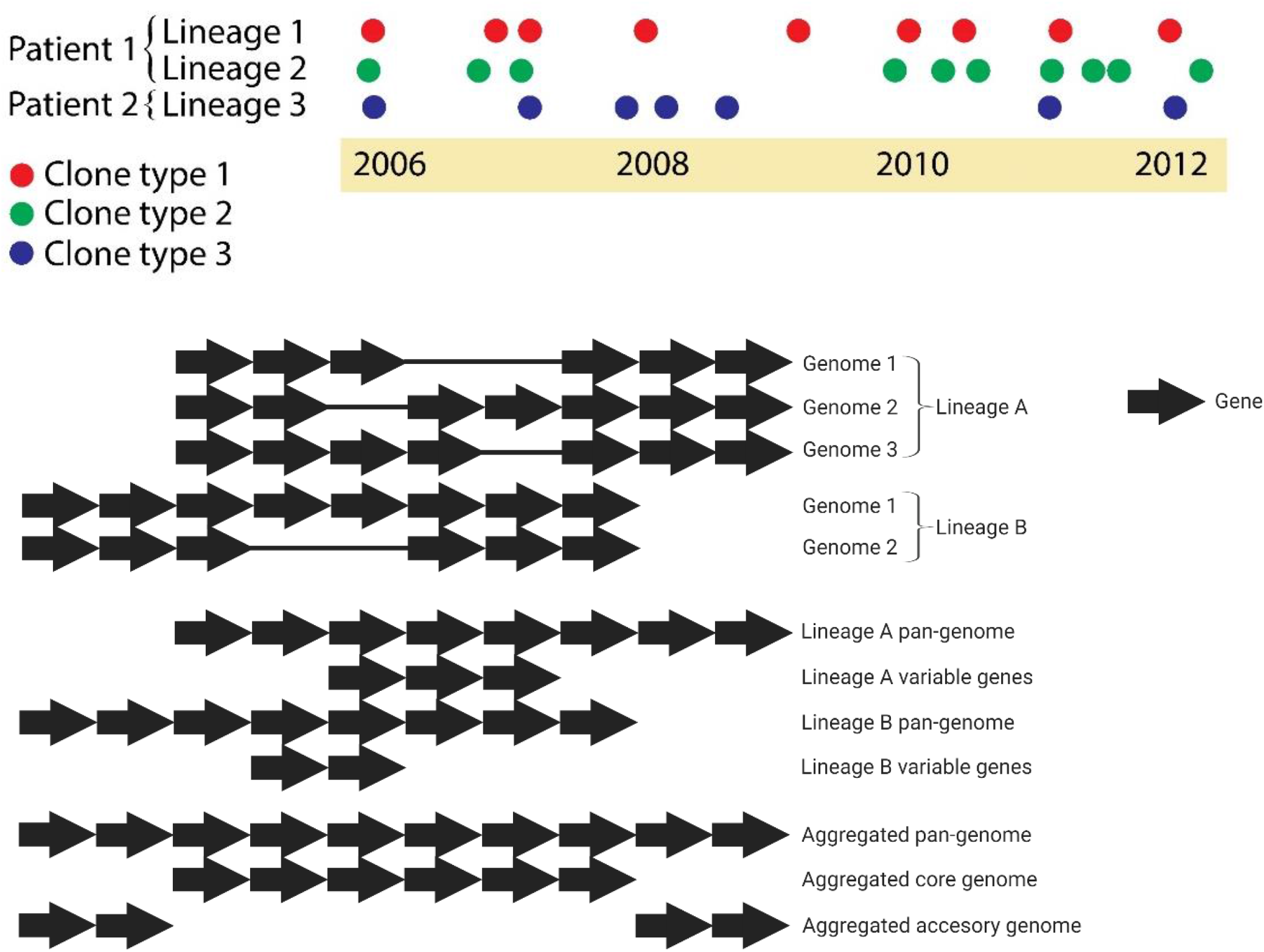
Schematic visualization of how bacterial lineages, lineage pan-genomes, within-lineage variable genes, and aggregated pan-, core- and accessory genome were defined in this study.

### Pan-genomes and identification of gene presence-absence

We analyzed the 423 genomes in a two-step process to identify genes that show variation within and between lineages, respectively. First, we compared the genomes of isolates of the same lineage to determine the full set of non-redundant genes found within the lineage, i.e. the lineage pan-genome. The lineage pan-genome consisted of (a) genes present in all isolates of the respective lineage, i.e. the lineage core genome, and (b) genes present in only some of the lineage isolates, i.e. genes that have been lost or acquired during the infection and here referred to as variable genes (Figure 1). Lineage pan-genomes consisted of 5,607–7,008 genes longer than 150 bp of which 0–473 were variable genes (median 44 variable genes).

Second, we compared the lineage pan-genomes to determine the full set of 14,462 non-redundant genes found across all lineages, i.e. the aggregated pan-genome (Figure 1 and Figure 2). The aggregated pan-genome consisted of a core genome of 4,887 genes, i.e. genes present in all lineage pan-genomes, and an accessory genome of 9,575 genes, i.e. genes present in only one or some lineage pan-genomes (Figure 1 and Figure 2). About half (4,932 genes) of the accessory genes were only present in a single lineage, and overall the lineage pan-genomes contained 0–540 (median 78) of such lineage-specific genes (Table S1). Furthermore, we found that all 335 genes reported to be essential genes in PAO1 and UCBPP-PA14 [28], were in the core genome.

**Figure 2.**
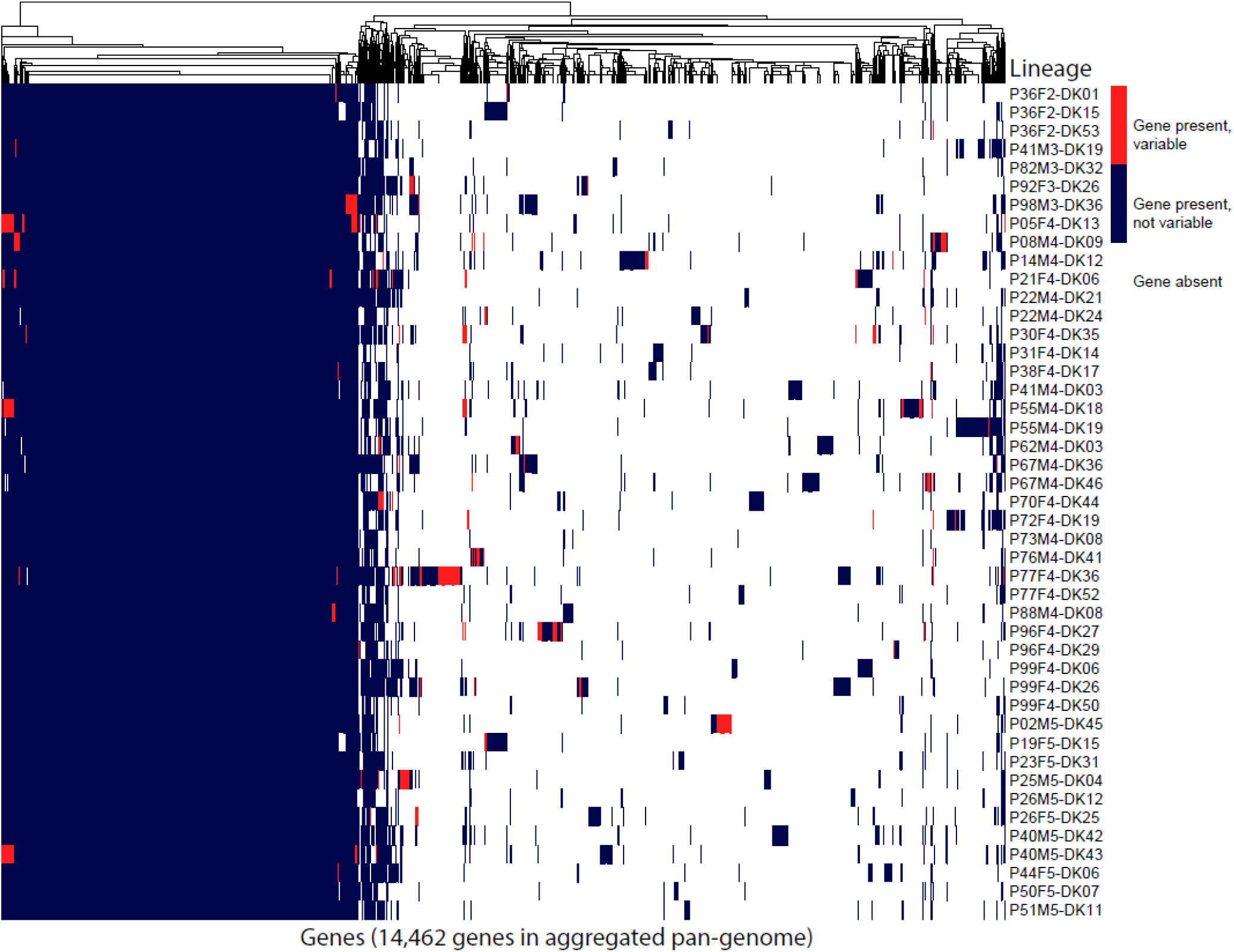
The heatmap of the aggregated pan-genome of 45 lineages containing all lineages with gene presence, absence and variability information.

We found that accessory genes were 16-fold more often variable within lineages (within hosts) than genes in the core genome (Supplementary materials, Table S1). While several factors might drive the higher turnover of accessory genes, one explanatory factor could be that the accessory genome has a higher amount of mobile genetic elements, such as prophage origin sequences. We used ACLAME database to identify and annotate phage and prophage sequences (longer than 150bp) in the core and accessory genomes, respectively. The accessory genome contained 116-fold more prophage genes, and these genes were highly variable over the course of infection: 58% of the prophage sequences in the accessory genome were variable within lineages.

### Changes in gene content in lineages over the course of infection

Next, we asked if the variable genes were either lost or acquired in bacterial linages. For this, we defined a gene as lost if it was present in the first isolate but absent in one or more of the later isolates, and a gene was defined as acquired if it was absent in the first isolate but present in one or more of the later isolates. Note, this definition of gene loss/acquisition might not be true as the first isolates may not represent the most recent common ancestor for the lineage. We found that the variable genes were most often likely lost. Out of 3,981 variable genes, 3,427 were present in the first isolates and absent in the later ones and the opposite was true for only 553 genes. Accordingly, we concluded that gene loss occurs at least 6 times more often than gene acquisition (Supplementary materials, Table S2).

Prophage sequences and plasmids are known to be mobile elements in bacterial genomes. Prophage genes were found in all 45 lineages by using ACLAME database. In 22 of the lineages, prophage genes were among the variable genes, and in 70% of cases (Supplementary materials, Table S2) the prophage genes were lost, i.e. they were present in the early isolates and absent in the later ones. Contrary, plasmid genes were not identified to be lost or acquired in any lineage (PlasmidFinder database was used to define plasmid genes).

257 genes in the aggregated pan-genome were related to virulence as per defined by VFDB database. Of these, 17 genes were variable in at least one lineage, and in 8 out of 17 cases, these genes were variable in more than one lineage. A two-sided Fisher’s exact test showed that virulence genes were in general less often lost/acquired than other genes (p-value = 4.45·10^−9^).

Seven genes in the aggregated pan-genome were related to resistance as per defined by ResFinder database. None of the resistance genes were lost/acquired in any lineages. Each isolate had 5–7 antibiotic resistance genes identified from ResFinder database (Supplementary materials, Table S1).

Out of 52 pathoadaptive genes reported by Marvig *et al.* (2015) [10], 9 were lost/acquired in lineages. No significant difference in lost/acquisition between pathoadaptive and non-pathoadaptive genes was found by performing a two-sided Fisher’s exact test (p-value 0.861).

Finally, we identified that genes are 21-fold more often lost or acquired in a group than individually (see example in Figure 3, Supplementary materials, Table S2), i.e. the loss/acquisition of 3,806 out of 3,981 variable genes congregated with the loss/acquisition of other genes while only 175 genes were lost/acquired alone.

**Figure 3.**
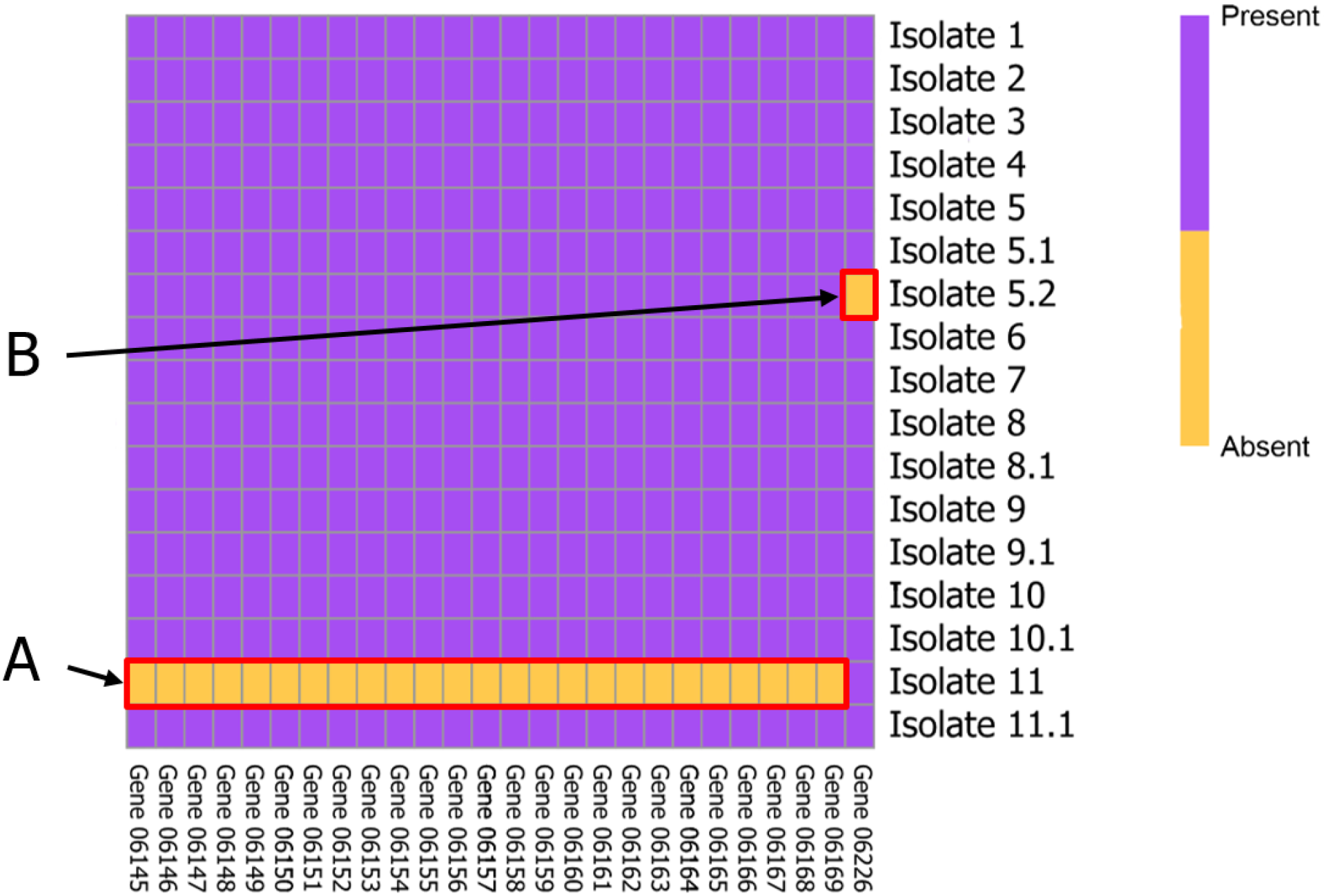
Heatmap that shows variable genes within lineage P98M3-DK36. Isolates are numbered according to date of sample. Earliest isolate is Isolate 1. (A) Multiple gene deletion during one deletion event and (B) individual gene deletion.

### Convergent evolution: same genes variable across lineages

While variable genes made a small fraction of the aggregated pan-genome, and the most variable genes were only lost/acquired in one lineage, some genes were observed to be variable in multiple lineages.

We defined genes as highly variable if they were identified as variable in ≥4 lineages. To ensure that the high variation was not due to technical artifacts of analysis, all highly variable genes were manually checked: (1) *P. aeruginosa* origin genes were mapped to a PAO1 reference genome and the coverage of the gene alignments was manually assessed, (2) for other genes the aggregated pan-genome was blasted by using BLAST+ suite [29] against isolate genomes and then the alignments were manually assessed. Out of initially 54 genes identified as variable in ≥4 lineages, 2 were removed after the manual check (detail explanation available in Materials and Methods section). Out of the 52 manually confirmed variable genes: 47 genes were variable in 4 lineages, 4 genes were variable in 5 lineages, and 1 gene was variable in 10 lineages (Figure 4).

**Figure 4.**
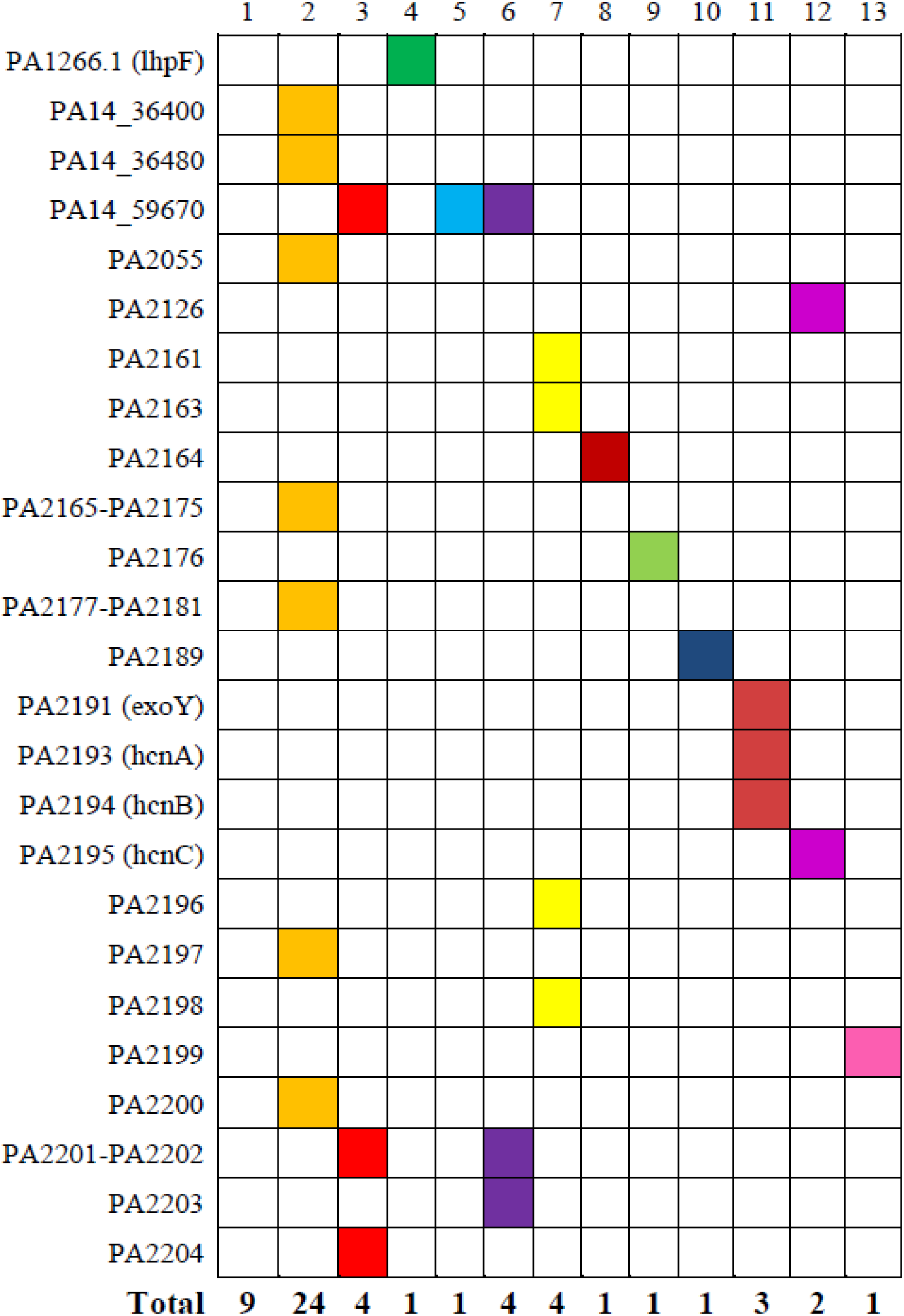
List of most variable genes and their function according to PseudoCAP annotation. The genes code for the following proteins: 1 – non-*Pseudomonas* origin (individual gene names not shown), 2 – hypothetical, unclassified, unknown, 3 – membrane proteins coding, 4 – amino acid biosynthesis and metabolism related, 5 – antibiotic resistance and susceptibility, 6 – transport of small molecules, 7 – putative enzymes, 8 – energy metabolism, 9 – two-component regulatory systems, 10 – Secreted factors (toxins, enzymes, alginate), 11 – central intermediary metabolism, 12 – transcriptional regulators, 13 – nucleotide biosynthesis and metabolism.

We annotated the highly variable genes according to PseudoCAP functional classes [30] (if present in PAO1/UCBBP-PA14 reference genomes) or by BLAST search against National Center for Biotechnology (NCBI) nucleotide collection (nr/nt) database. Most of the highly variable genes (34 of 52) were genes with hypothetical function or genes of non-*Pseudomonas* origin. The second largest groups of highly variable genes encoded membrane proteins (4 genes). Since genes encoding membrane proteins make up around 10% of the *P. aeruginosa* genome, we tested using a Fisher’s exact test if membrane genes encoding membrane proteins are more variable than expected when accounting for their abundance, and we concluded that it is not the case (p-value = 1.00). Other highly variable genes encoded proteins involved in amino acid and nucleotide biosynthesis, antibiotic resistance, transport, secreted factors, transcriptional regulation, and metabolism (Figure 4).

### *Convergent evolution of locus with hcnABC* and *exoY* genes

We identified that a group of 34 genes was lost/acquired in four lineages (P21F4-DK06, P05F4-DK13, P55M4-DK18 and P40M5-DK43). The 34 genes were orthologous of genes PA2161–PA2181 and PA2189-PA2204 in reference genome PAO1. We noted that three of the lineages (P05F4-DK13, P55M4-DK18 and P40M5-DK43) did not have genes PA2182–PA2188 (genes flanked by PA2161–PA2181 and PA2189-PA2204 in PAO1 reference genome) in their lineage pan-genomes, and that genes PA2182– PA2188 were variable and congregated with genes PA2161–PA2181 and PA2189-PA2204 in the fourth lineage (P21F4-DK06), so we concluded that the 34 genes were likely lost/acquired together rather than in separate two events (Figure 5 shows the genetic region of the group of 34 variable genes). Further, we aligned reads from each of the lineages to PAO1 reference genome to show that parallel deletion/insertion of the 34 genes was the result of larger yet different deletions/insertions in the individual lineages (i.e. the 34 genes were the shared overlap among four different deletions/insertions in the same genomic region; see Supplementary materials, Figure S1 and Figure 5).

**Figure 5.**
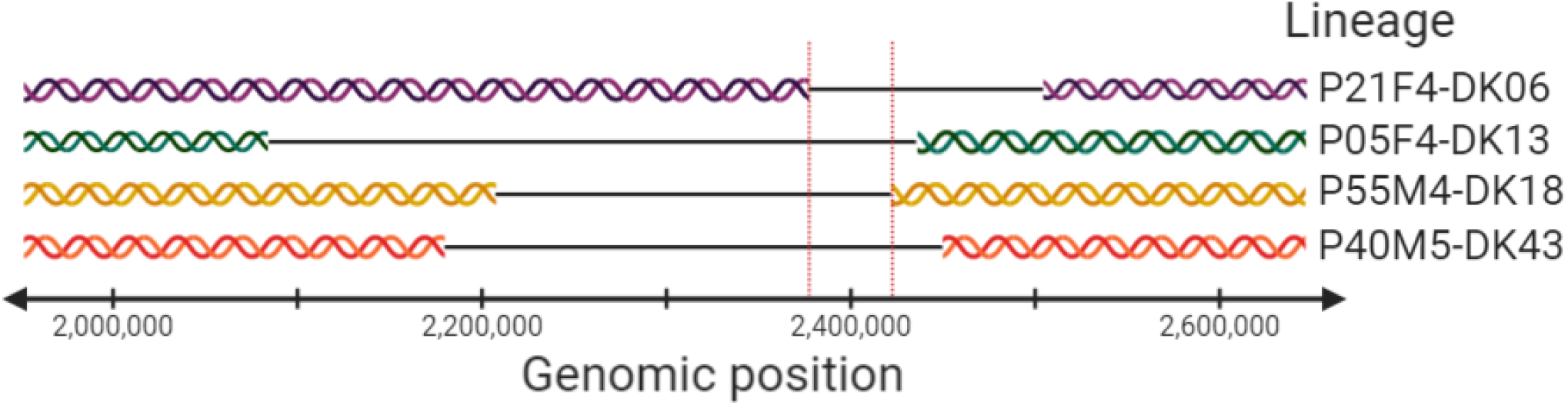
Genetic regions of four lineages where the same group of 34 genes is lost. Red vertical lines show the overlapping genetic region lost in all four lineages.

In 3 out of 4 lineages, genes were present in early isolates and absent in late isolates. For lineage P40M5-DK43, the 34 genes were present in the later of only two lineage isolates that were sampled less than a year apart, so it is likely that two isolates represent different sublineages where one sublineage did not lose the genes while another one did. Also, genes PA2161–PA2204 were present in all 45-lineage pan-genomes, suggesting that genes PA2161–PA2204 were also present in ancestor of lineage P40M5-DK43 and thus were lost during course of infection.

17 of the 34 genes were annotated as ‘Hypothetical, unknown or unclassified’ by PseudoCAP, other genes were coding for ‘Putative enzymes’ (4 genes), ‘Transport of small molecules’, ‘Membrane proteins’, ‘Central intermediary metabolism’ (3 genes) or ‘Energy metabolism’, ‘Two-component regulatory systems’, ‘Secreted factors’, ‘Transcriptional regulators’ and ‘Nucleotide biosynthesis and metabolism’ (1 genes) (Supplementary materials, Table S4). Some of these proteins have more than one function assigned by PseudoCAP. Literature search identified that four of the genes (*hcnABC* and *exoY* encoding hydrogen cyanide synthase and type III secreted protein, respectively) are known to play a role in virulence and pathogenesis of *P. aeruginosa.* [30, 31]

### Covergent evolution in prophage related genes and genomic islands

Six of the 52 highly variable genes were identified as prophage origin genes originating from different *P. aeruginosa* prophages similar to the genes from phi1 and phi2 *Pseudomonas* phages. The variable groups of 56–82 genes which included the 6 most variable prophage genes might as well be yet undescribed genomic islands as they are adjacent to tRNA encoding genes, contain both prophage origin and *P. aeruginosa* origin sequences, and are longer than 10,000 bp.

As the mobility of genomic islands could explain high variability of these gene regions, we predicted GIs with IslandViewer 4. All six prophage origin genes were predicted to be part of GIs, e.g, exemplified by a 78 gene deletion in a genomic island encoding virulence factors in lineage P67M4-DK46 (Figure 6). While IslandViewer4 predicted on average 40 GIs (range 15–59) per lineage; we note that, as the analyzed genomes were not complete (i.e. in scaffolds), the GI prediction should be interpreted carefully. E.g. GIs were often predicted at the ends of scaffolds.

**Figure 6.**
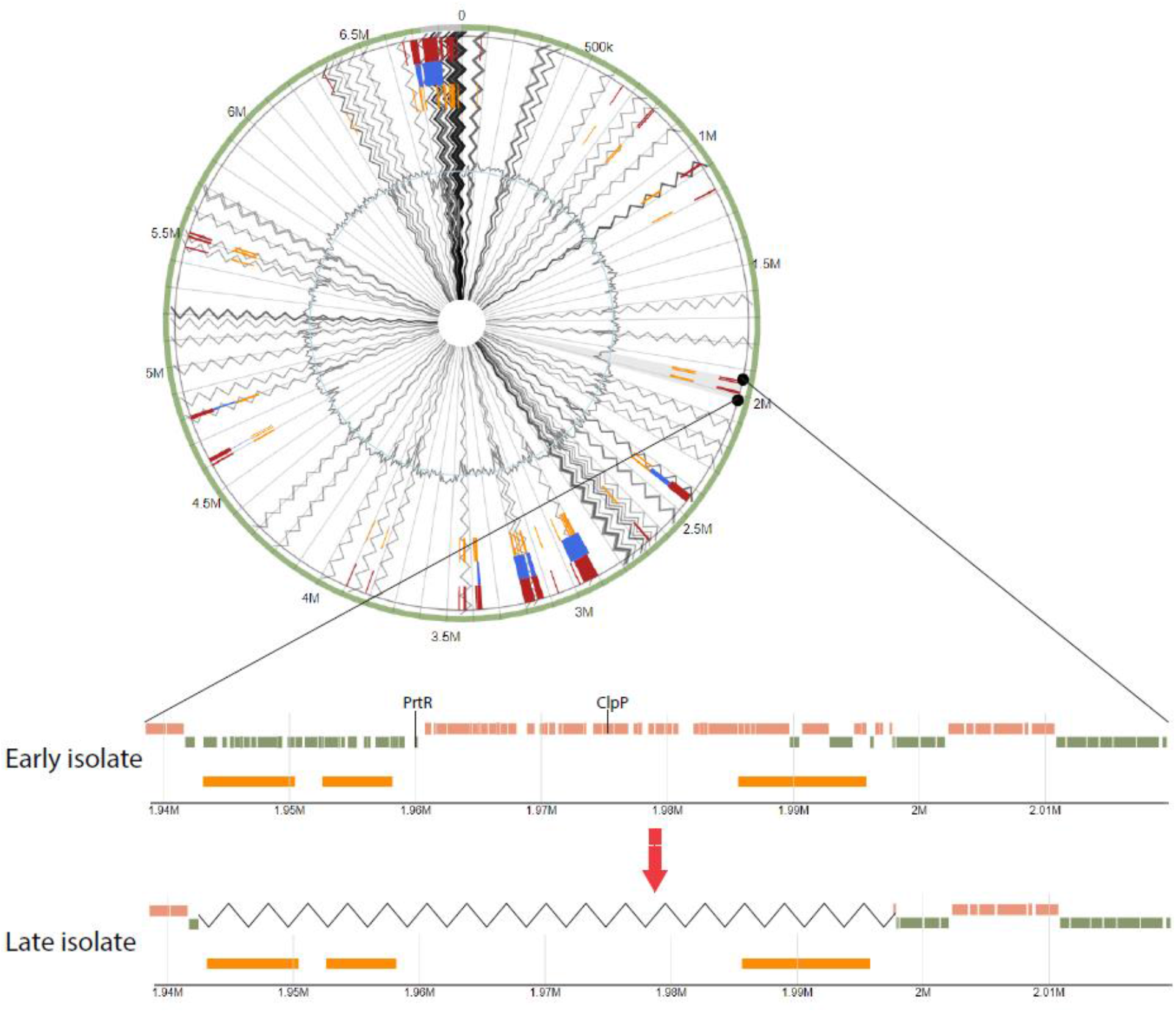
Genomic island predictions for an early and late isolate of lineage P67M4-DK46, respectively. Zoomed region shows gene loss (zigzag line) in the late isolate. Possible virulence factors in the zoomed region are marked in the early isolate. Orange blocks in early and late isolates show predicted genomic islands.

On average 90% (87–92%) of genes in the predicted GIs code for hypothetical proteins. Therefore, it is difficult to define the function of most of the genes present in GIs. However, possible drivers of the loss of predicted GIs were identified: we identified homologs of genes coding for Clp protease (7 lineages) or *prtR* gene (5 lineages) as parts of predicted GIs which are known to be related to bacterial virulence and pathogenicity. [32, 33] In addition, other probable virulence factors were identified in multiple lineages (Supplementary materials, Table S3).

### Overall population structure and SNP and gene distances

We wanted to understand our results on lineage genomes in the context of the overall population structure of *P. aeruginosa*. Accordingly, we determined the genetic relationship of the 446 isolates based on either SNPs in the core genome or gene presence-absence. Both the SNP- and the gene-based phylogeny clustered the isolates according to clone type and patient (Figure 7). Also, both phylogenies showed that lineages clustered into one of two overall groups with either reference strain PAO1 or UCBPP-PA14, respectively. Core genome pairwise SNP distance between lineages was on average 31,909 (22–67,325) SNPs while the gene content difference was on average 1,142 (13–2,250) genes (one random isolate was chosen to represent each lineage to avoid overrepresentation of some lineages and underrepresentation of others, Figure 8). Moreover, the average between lineage diversity was 19,853 (22–24,957) SNPs and 1,043 (28–2,250) gene differences in PAO1 group, respectively, while the diversity within UCBPP-PA14 group was 35,191 (44–67,325) SNPs and 1,217 (13–1,775) gene differences, respectively. After performing Wilcoxon’s rank sum test on the distributions in the two groups, a significant difference between the groups was identified with p-value <2.2·10^−16^ for pairwise SNP distance and p-value=2.57·10^−13^ for pairwise gene difference distance showing that the variability of SNPs and genes is lower within the PAO1 group than within UCBPP-PA14 group.

**Figure 7.**
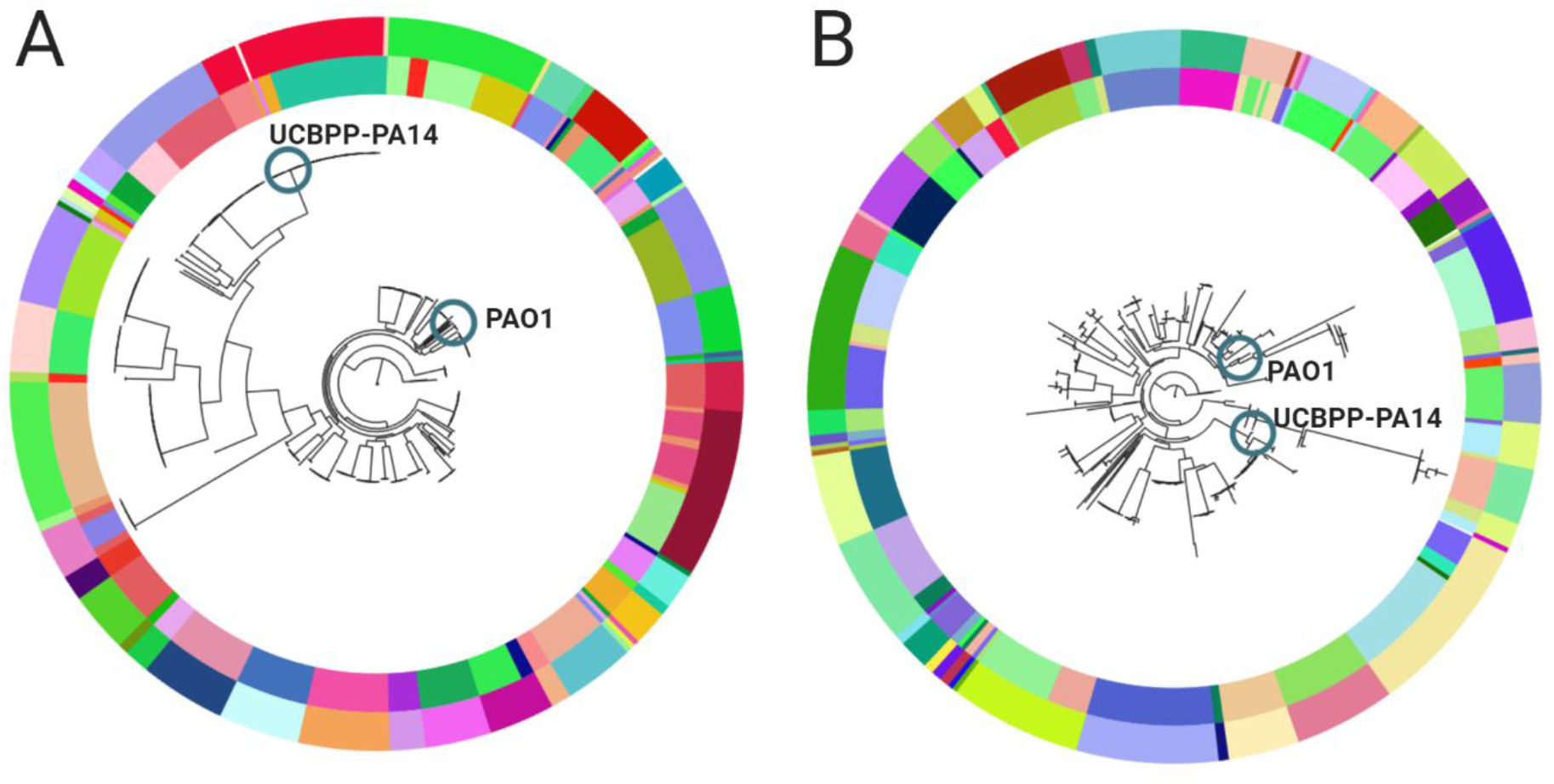
Phylogenetic trees of 446 *P. aeruginosa* samples (A) based on core genome SNPs and (B) based on gene presence-absence. The outer color circle of the trees denotes clone type, and the inner circle denote patient. Blue circles denote the position of reference genomes.

**Figure 8.**
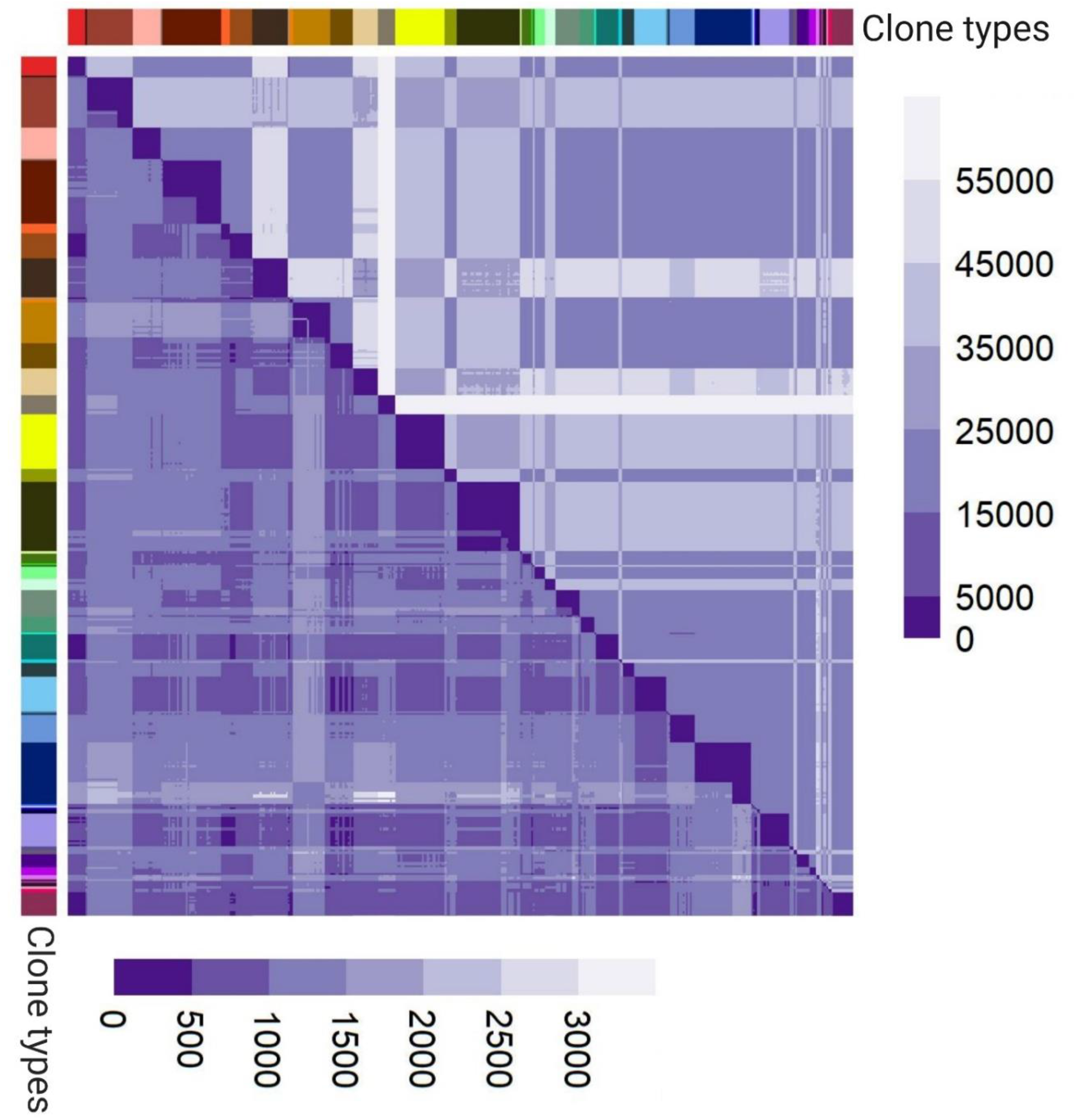
Heatmap of pairwise SNP distances (top triangle) and gene distances (bottom triangle) between *P. aeruginosa* isolates with clone type annotation on the left side and on top.

### Core genome SNP distances to define clone types

Isolate clone types were previously defined by Marvig *et al.* (2015) [10] by one-to-one pairwise alignment of genomes, and genomes that differed by >10,000 SNPs were defined as of different clone types. This way of defining clone types was ambiguous as the number of SNPs was determined on the basis on alignment of genes (sequences) shared between only the two respective genomes rather than based on a fixed set of genes. Therefore, we employed the identified core genome in a more robust method to define clone types and using the core genome as a fixed basis for determining SNP distances, we found that 5,000 SNPs clearly confirmed and defined clone types (Supplementary Materials, Figure S2). Furthermore, we developed the method into a workflow—Pactyper (https://github.com/MigleSur/Pactyper)—that can be used to define clone types from the core genome and sequence read data of any collection of bacterial isolate.

Finally, we found that the ratio of core genome SNPs per difference in gene content was on average 3.5 (0.01–115.00, median: 0.66) and 30.2 (7.22–88.00, median: 28.95) within and between clone types, respectively. Using a Wilcoxon’s rank sum test, we concluded that the difference between two groups is statistically significant with p-value<2.2·10^−16^.

### Re-analysis of other study to compare size of pan- and core genome

We analyzed public available genome sequencing data on a collection of 1,139 isolates that have previously been included in a pan- and core genome analysis by Freschi *et al.* (2019) [19], to compare the pan- and core genome sizes between different collections of isolates. When we used the same method as for our own isolate collection (GenAPI), we found that the 1,139 isolates from Freschi *et al.* shared a core genome of 2,360 genes within a pan-genome of 38,017 genes. When we used the method of Freschi *et al.* (2019) (SaturnV), the core and pan-genome consisted of 619 and 43,703 genes, respectively. If the core genome was defined as all genes present in at least 99% of the samples, the core genome consisted of 4,870 and 3,879 genes for GenAPI and SaturnV, respectively.

## Discussion

By analysis of genome sequences from 45 longitudinally sampled *P. aeruginosa* lineages from CF patients, we determined the micro-evolutionary dynamics of gene loss and acquisition in lineages of bacteria evolving in a human host environment. While similar analyses of within-host bacterial evolution have investigated within-host gene loss and acquisition, our collection enables comparative analysis across multiple genotypically different strains (45 lineages distributed on 34 clone types) of the same species. Hereby, we not only identify events of gene loss or acquisition in the individual lineages, but also analyze this in the context of the gene variation across lineages, *i.e*. in the context of the species pan- and core genome.

### Pan- and core genomes

The aggregated pan-genome across lineages had 14,743 genes in total, and 4,887 genes were present in all 45 individual lineage pan-genomes (*i.e.* here defined as the core genome across lineages). Our findings are similar to study by Hilker *et al.* (2015) [34] that found the genomes of 21 *P. aeruginosa* strains to share a core of 4,748 genes of a pan-genome comprising 13,527 genes. In contrast, our pan- and core genome sizes are significantly different from recent findings by Freschi *et al*. (2019) [19] where the pan- and core genomes were 54,272 and 665 genes, respectively. By re-analysis of the dataset from Freschi *et al.* with both our (GenAPI) and original (SaturnV) method, we show that the difference in pan- and core genome sizes can largely be ascribed to differences in bioinformatics analyses as pan- and core genome sizes from Freschi *et al.* converge towards findings from this and other studies [34, 35, 36, 37] when we re-analyzed the data with GenAPI. We have previously compared tools SaturnV and GenAPI [15], and we suggest that the relatively small size of the core genome reported by Freschi *et al.* (665 genes) is due to false negative calls on genes with incomplete assemblies [38].

### Within host gene loss and acquisition

We found that genes were six times more often lost than acquired within lineages during within host evolution. It remains unknown if the lack of gene acquisition is limited due to availability of donor DNA, mechanisms of DNA uptake, or selection (either selection against acquisition of genes or lack of selection for acquisition of genes). Nonetheless, we find our results to be in line with previous hypotheses that genomes are selectively reduced during the course of infection [39, 40].

Most of the genes that were lost or acquired within the host, were part of the genome that was not shared across lineages (*i.e.* the accessory genome). The relative low turnover in the core genome of 4,887genes shared by all lineages suggests that, while these genes are not essential *per se*, they may be generally important for survival in the conditions met by *P. aeruginosa* in human host environment. Accordingly, they are maintained in the genomes.

In contrast, prophage genes were the genes that were most often lost or acquired within the host, and also prophages were putatively the major source of new genetic material. 268 of 462 (58%) of prophage genes in the aggregated pan-genome were variable within hosts, and despite taking up only 3% of the aggregated pan-genome, prophage genes constituted 9.4% of all within-host variable genes. This confirms that prophage-facilitated gene flux is abundant and supports the conclusions from other studies that prophages play an important role in *P. aeruginosa* CF infections [41, 42].

### Convergent evolution and adaptive loss of virulence

The sampling from multiple lineages (and across multiple patients) offered the opportunity to detect genes that were loss or acquired in independently in parallel evolving lineages. While most genes were only variable in a single lineage, we found 52 genes to be variable within ≥4 lineages. The observed parallel loss or acquisition of the same genes across lineages may be driven by selection for loss and acquisition of certain genes in the host environment. It has previously hypothesized that virulence factors are selected against in CF infections, and in agreement with this we found that 34 of the 52 highly variable genes were lost as part of a genomic region encoding virulence factors hydrogen cyanide synthase (*hcnABC*) and type III secreted protein ExoY (*exoY*). It has also been shown by Wee *et al.* (2018) [31] that selective pressures for losing *hcnA*, *hcnB*, *hcnC* and *exoY* genes exist, and we notice that Wee *et al.* also observed deletions of varying sizes around the respective genes. Furthermore, the selective pressure for loss of *hcnABC* locus virulence genes was recently shown to possibly be related to increased antibiotic resistance in multidrug resistant strains. [40]

We notice that while virulence genes *hcnABC* and *exoY* are among the most variable genes, we found that virulence genes in general were less often variable within lineages. This may be counter-intuitive if loss of virulence is beneficial for bacteria in chronic infections; nonetheless, we recognize that virulence factors may instead be downregulated rather than deleted as suggested by Rau *et al.* (2010). [41]

Genes were 21 time more often observed to be lost or acquired as groups of genes as opposed to as single-gene losses or acquisitions. This observation is in line with previous studies [39, 40] and illustrates how the presence of specific genetic elements enables and defines mobilization of genes. This is illustrated by six of the 52 highly variable genes are prophage genes, and prophage regions often act as mobile elements. Accordingly, these six prophage genes were part of groups of 56–82 genes that were deleted together and constituted genomic islands.

All gene groups that were lost with the six highly variable prophage genes contained the genes coding for Clp protease and *prtR* gene. PrtR is required for type III secretion system [33] and Clp protease is inducing virulence by regulation of flagella gene expression and ultimately increasing bacteria adhesion. [32] Accordingly, the frequent loss of *prtR* and Clp protease genes adds to our observation that virulence factors are lost during infection. This loss of virulence may be positively selected in the host environments as the virulence factors activates the host immune response; hence loss of virulence helps the bacteria to hide from the immune defense. Two of eight lineages with loss of Clp and *ptrR* also lost *hcnABC* loci and *exoY* genes, and as such we observed no evidence that loss of the different virulence factors was mutual exclusive or concurrent (Supplementary materials, Table S5).

### Clone types and population structure

We developed a tool, Pactyper, that meets the need for a standardized way to define clone types and quantify genome SNP distances at the population level. We described the population structure of our *P. aeruginosa* population of 446 isolates using both SNPs (Pactyper) and gene absence/presence (GenAPI) information, and in both ways we identify two major phylogenetic clusters: one with PAO1 and one with UCBPP-PA14, which agree with previous studies by Hilker *et al.* (2015) [34] and Stewart *et al.* (2014) [42]. Furthermore, we showed that the SNP and gene difference is significantly lower among PAO1-like isolates compared with UCBPP-PA14-like isolates. Finally, we defined that there is a significantly less SNPs per gene loss/acquisition in isolates belonging to the same clone type than in isolates from different clone types.

In summary, we used a genome-wide and hypothesis-free gene presence-absence analysis approach to identify the main patterns of *in vivo* bacterial micro-evolution. Furthermore, we used the defined core genome to develop a stable and discriminatory typing scheme for *P. aeruginosa* clone types (Pactyper, https://github.com/MigleSur/Pactyper). In sum, our analysis adds to the knowledge of how prevalent loss or acquisition of genes is within bacteria evolving in the human host environment and provides a basis to further understand how gene loss and acquisition plays a role in host adaptation.

## Author statements

### Authors and contribution

R.L.M., H.K.J, and S.M. conceived the study. R.L.M. and F.C.N supervised the study. M.G., R.L.M. and H.K.J. defined the methodology. M.G. and R.L.M. designed the bioinformatics workflows for the analysis. M.G. conducted the analysis. M.G. and R.L.M. analyzed and interpreted the data. M.G. prepared the manuscript draft and visualizations. R.L.M., H.K.J., F.C.N and S.M. reviewed and edited the draft.

## Acknowledgments

We thank Anders Krogh for valuable discussions throughout the study. Also, Figure 5 and Figure 8 were created with BioRender (https://app.biorender.com/).

